# Lévy statistics define anxiety and depression in mice subjected to chronic stress

**DOI:** 10.1101/2023.06.18.545232

**Authors:** Qinxi Li, Yuru Nie, Xiaojie Li, Yiping Luo, Bangcheng Zhao, Ni Zhang, Weihong Kuang, Chao Tian, Daojun Chen, Yingqian Zhang, Zhe Wu, Zhihui Zhong

**Author notes:** Corresponding Author 1: Zhihui Zhong Laboratory of Neurological Disease Modeling and Translational Research West China Hospital, Sichuan University No.37 GuoxueAlley, Wuhou District Chengdu, Sichuan, 610041, China; Corresponding Author 2: Zhe Wu School of Life Science and Technology University of Electronic Science and Technology of China 2006 Xiyuan Avenue, High-tech Zone (West District) Chengdu City, 611731, China Corresponding Author 3: Yingqian Zhang Laboratory of Neurological Disease Modeling and Translational Research West China Hospital, Sichuan University No.37 GuoxueAlley, Wuhou District Chengdu, Sichuan, 610041, China. These authors contributed equally to this work (Qinxi Li, Yuru Nie, Xiaojie Li).

## Abstract

**Introduction:** Anxiety and depression are recognized as adaptive responses to external stressors in organisms. Current methods for evaluating anxiety and depression in rodents are both burdensome and stressful. The objective of this investigation is to explore a simplified methodology for identifying stress-induced and stress-free states, as well as anxiety and depression levels, by analyzing the movement patterns of rodents.

**Methods:** To address this issue, we utilized Lévy statistics to examine the movement patterns of stressed rodents and compared them to non-stressed controls. We employed the two-dimensional Kolmogorov-Smirnov test to identify significant differences in the γ and μ parameters derived from Lévy flight (LF) between anxiety, depression, and control mice. Additionally, we employed the support vector machine algorithm to optimize the classification of each group.

**Results:** Our analysis revealed that stressed mice displayed heavy-tailed distributions of movement velocity in open fields, resembling the movement patterns observed in animal predators searching for scarce food sources in nature. In contrast, non-stressed mice exhibited a normal distribution of speed. Notably, the effectiveness of this methodology in the field of drug discovery was confirmed by the response of stressed mice to fluoxetine, a well-established selective serotonin reuptake inhibitor (SSRI).

**Conclusion:** This study unveils a previously unidentified statistical walking pattern in mice experiencing anxiety and depression. These findings offer a novel and accessible approach for distinguishing between anxiety, depression, and healthy mice. This method provides a one-step gentle approach (free walk in an open field) instead of the traditional multi-step stressful tests.

## Introduction

In 2019, the World Health Organization reported that one out of every eight individuals worldwide lives with one or more mental disorders ^1^. The COVID-19 pandemic resulted in a significant increase in depression and anxiety cases within a year ^2^. Chronic stress is the principal environmental factor that triggers a decline in mood, leading to depression and anxiety ^3^.

To mimic these disorders, rodent models of stress, particularly the chronic unpredictable mild stress (CUMS) model, have been extensively employed since the early 1980 ^4^. In CUMS and its derived models, mice or rats are subjected to constant but unpredictable mild stressors, resulting in the development of depression-like or anxiety-like behaviors, mimicking the core symptoms of clinical depression and anxiety, such as anhedonia and acquired helplessness ^5^. CUMS has demonstrated its excellent reliability and validity in the realm of drug discovery ^6^.

Rodents’ response to stress is typically observed through sequential behaviors, including anhedonia, abnormal weight, and poor coat condition. In order to quantitatively assess the severity of mental disorders and distinguish between anxiety and depression-like behaviors in experimental animals, conventional neurobehavioral studies such as the elevated plus maze (EPM) and light/dark box (LDB) test for anxiety ^6,7^. The forced swim test (FST) and tail suspension test (TST) for depression are commonly employed ^8,9^. The open field test (OFT) serves as a valuable tool for assessing both anxiety-like and depressive-like behaviors in mice. In terms of anxiety, it is characterized by increased distance traveled and prolonged rearing exploration. On the other hand, depressive-like behavior is indicated by an extended duration spent in the corners of the open field and an avoidance of crossing the central zone^10^. Nevertheless, it is important to acknowledge that each of these established methodologies has its own set of limitations. For instance, Cheryl D. Conrad has pointed out that the assessment of anxiety through the OFT does not necessarily correlate with anxiety assessed using the EPM ^11^. Additionally, the TST and FST assays only capture a single facet of depression ^9^, while the EPM may induce anxiety in animals during testing, thereby complicating the evaluation process ^12^. Furthermore, the very procedures involved in these experiments impose additional stress on the animals ^13^, which can interfere with the interpretation of data and restrict the translational relevance of the studies.

Lévy flight (LF), named after the esteemed French mathematician Paul Levy, is a “random walk” initially denoted a stochastic traversal characterized by a heavy-tailed probability distribution of step lengths. Within mathematics, the “random walk” is an algorithm that takes cues from nature, involving a sequential progression of successive steps with random distances or velocities ^14,15^. In the natural world, the movement of animals exhibits different patterns based on the availability of food. When resources are plentiful, animal trajectories tend to resemble “Brownian motion”, characterized by frequent revisits to previously explored areas, a phenomenon often termed “oversampling” ^16^. In contrast, animals opt for more extensive movements when food and resources become scarce, manifesting as “heavy tails” in the probability distribution of step lengths. In statistical terms, “heavy tails” describes the phenomenon wherein the tails of a probability distribution exhibit slower decay. This implies that extreme events or outliers are more likely to occur than distributions with lighter tails^17^. The “LF foraging hypothesis” suggests that this motion pattern enhances their chances of survival ^15^. The applications of LF encompass a broad spectrum, including the analysis of earthquake data, financial mathematics, cryptography, signal analysis, and a wide array of practical applications in the fields of astronomy, biology, and physics ^18,19^.

The hypothesis in this study posits that laboratory animals would transition from “Brownian” to “LF-like” motion patterns due to evolutionary pressure encoded in their genetic makeup. In this study, our objective is to examine the potential of “LF-like” motion patterns in distinguishing between stressed and non-stressed mice and further evaluate their effectiveness in differentiating anxiety and depression mice within the stressed group. These methodologies possess practical implications in the development of a streamlined single-step experimental technique for quantifying the severity of anxiety and depression induced by stress. Moreover, this approach holds the potential to minimize the extra stress encountered by animals during repetitive experimental protocols while concurrently enhancing the efficacy of drug discovery endeavors.

## Results

### Mice exposed to CUMS displayed characteristic anxiety-like or depression-like behaviors

Using a modified CUMS method (Figure 1A), we successfully induced anxiety or depression in mice models, which were validated using industry standards (Figure 1). To exclude subjects with naturally low activity levels, either physically or mentally, as described in the method section, 145 mice underwent an SP test (Figure supplement 1A) and an OFT (Figure supplement 1B). After two weeks of modeling, we utilized SP and OFT to assess the efficacy of CUMS induction. The control group excluded a preference rate of less than 80% for SP, whereas the CUMS group eliminated a preference rate higher than 80% (Figure supplement 1C). Seven mice were eliminated from the study on Day 14, as they did not enter the center of the OFT fewer than five times, following the exclusion criteria (Figure supplement 1D). The SP test was conducted thrice throughout the entire experiment, and the SP test conducted on the 42nd day, the CUMS + FXT sucrose preference percentage exhibited a significant increase when compared to the CUMS + saline group (Figure 1B). The mice subjected to CUMS for 42 days showed lower body weight. In the CUMS + FXT group, the body weight on the 42nd day exhibited a significant increase compared to the CUMS + saline group. (Figure 1C). On the 42nd day, the control group demonstrated a significantly elevated food intake in comparison to the CUMS + saline group (Figure 1D). In contrast, the control group exhibited a notably lower coat state score than that of the CUMS+saline group (Figure 1E). Nevertheless, no significant disparities were observed in the food intake and Coat state score between the CUMS+saline group and the CUMS+FXT group. During 37 days of treatment, the success of the CUMS model was validated by administering fluoxetine (FXT) to CUMS mice, with the observation that the CUMS + Saline mice showed fewer center entries and distance during OFT (Figure 1F-H). Neurobehavioral tests, including EMP and LDB, were conducted on all mice from day 44 to day 50 to evaluate anxiety-like behavior. Additionally, TST and FST were employed to assess depressive-like behavior. The CUMS + Saline group exhibited less exploration time in the open arm of EPM and light box in LDB (Figure 1I-N), as well as reduced struggling time in TST and FST, as shown in Figure 1 O-R. Conversely, FXT treatment increased the number of struggle times in the CUMS + FXT group.

**Figure 1.**
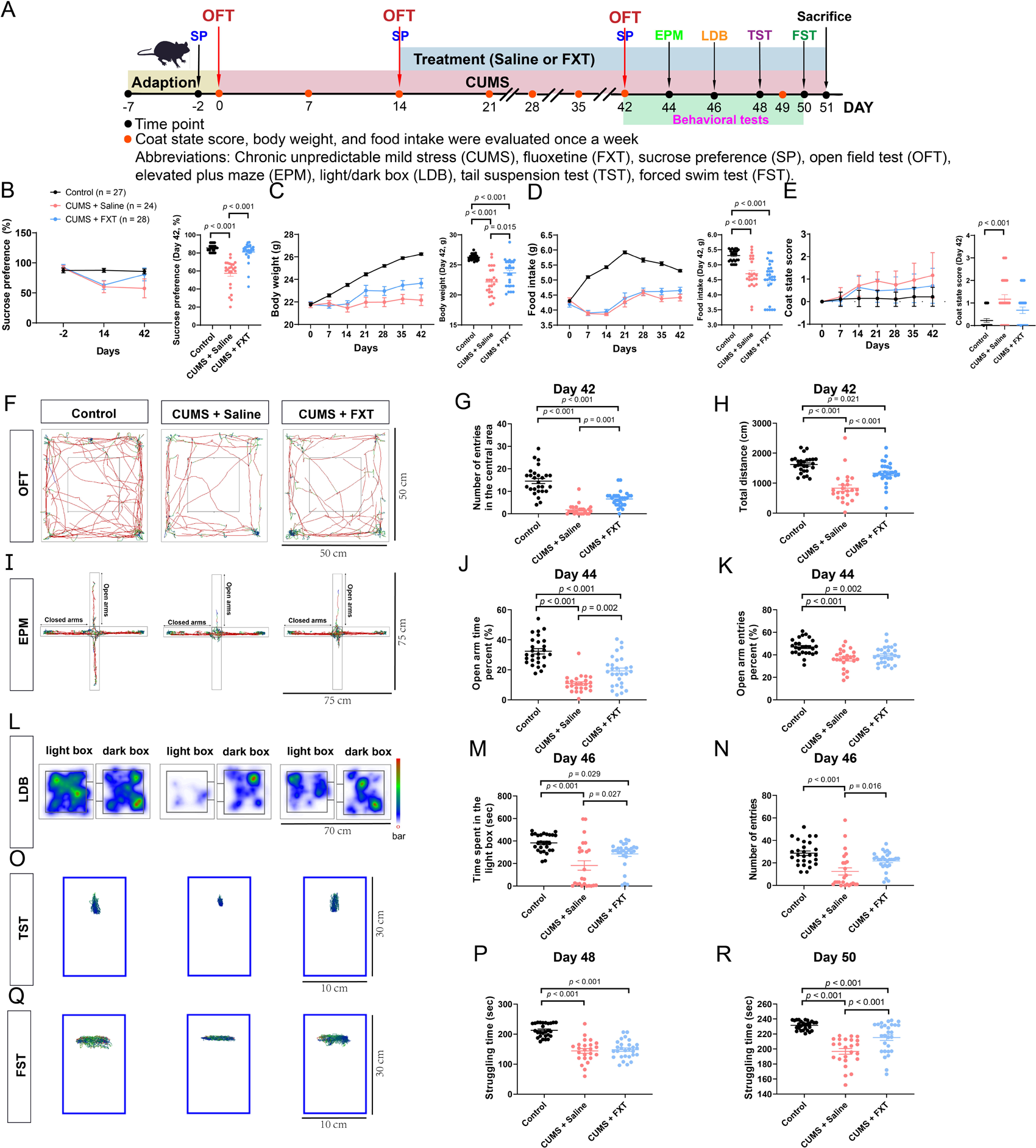
Timeline and validation of the experimental design for the CUMS Model. (**A**) CUMS experimental design demonstrated anxiety-like and depressed behaviors. Groups were assigned based on OFT results, with drug administration starting on day 14. (**B**) As depicted in the left Figure, exposure to the CUMS paradigm resulted in a substantial manifestation of anhedonia, as evidenced by the SP test. Additionally, on the 42nd day of the experiment, discernible differences in the percentage of water preference were observed among the various groups, as displayed in the right Figure. (**C-E**) In the left Figure, the CUMS paradigm was found to induce significant alterations in coat state score, body weight, and food intake. Furthermore, on the 42nd day of the experiment, the right Figure depicts a comparison of variations among the three groups. (**F**) On the 42nd day, OFT trajectories were recorded and analyzed. (**G**) On the 42nd day, mice in the CUMS + FXT group exhibited a higher number of center entries during the OFT. (**H**) CUMS + FXT mice exhibited reduced anxiety- and depression-like behaviors, as indicated by significantly greater distances traveled during the OFT. (**I-J**). During the EPM test, CUMS + Saline group spent less time exploring the open arm, with a notable difference observed between the CUMS + Saline and CUMS + FXT groups.(**K**) No significant difference in behavior was observed between the CUMS + Saline and CUMS + FXT groups. (**L-M**) CUMS + Saline group exhibited significantly reduced exploration time in the light box during the LDB test. (**N**) On the 46th day, the control group had a significantly greater entry during the LDB test. (**O-R**) During the TST and FST, mice in the CUMS + Saline group exhibited significantly more extended periods of immobility. The presented data represent mean ± s.e.m. Statistical significance was established when *p*-values were below 0.05. The number of animals is indicated on the graphs. A repeated measures analysis of variance (ANOVA) was employed to assess potential disparities among groups. In cases where interactions between an index and days were observed, we evaluated the distinctions among each group’s final time points, encompassing factors such as SP test outcomes, coat state scores, body weight, and food intake. In instances where no interactions were detected, subsequent post-hoc analysis, employing Bonferroni’s multiple comparison tests, was conducted.

### The velocity of mice walking in an OF showed a pattern of LF

Due to spatial confinement in limited arenas, rodent trajectories are often truncated by the boundaries, making statistical distributions a better model for walking speed than rodents’ walking trajectories to reveal mental states under stress. In this study, the walking rates of each mouse group were plotted against time in Figure 2A, and the frequency of walking speed was computed and represented as a bar graph in Figure 2B-C. Additionally, two probability density functions (PDFs), the LF distribution and normal distribution (Equation 3, 4), were fitted to the walking speed statistics for each mouse category. The goodness of Fit (R-good) quantitatively represented curve fitting quality. Results showed that control mice’s walking behavior followed normal and LF distributions, while CUMS mice had an LF distribution, as depicted in Figures 1D-E.

**Figure 2.**
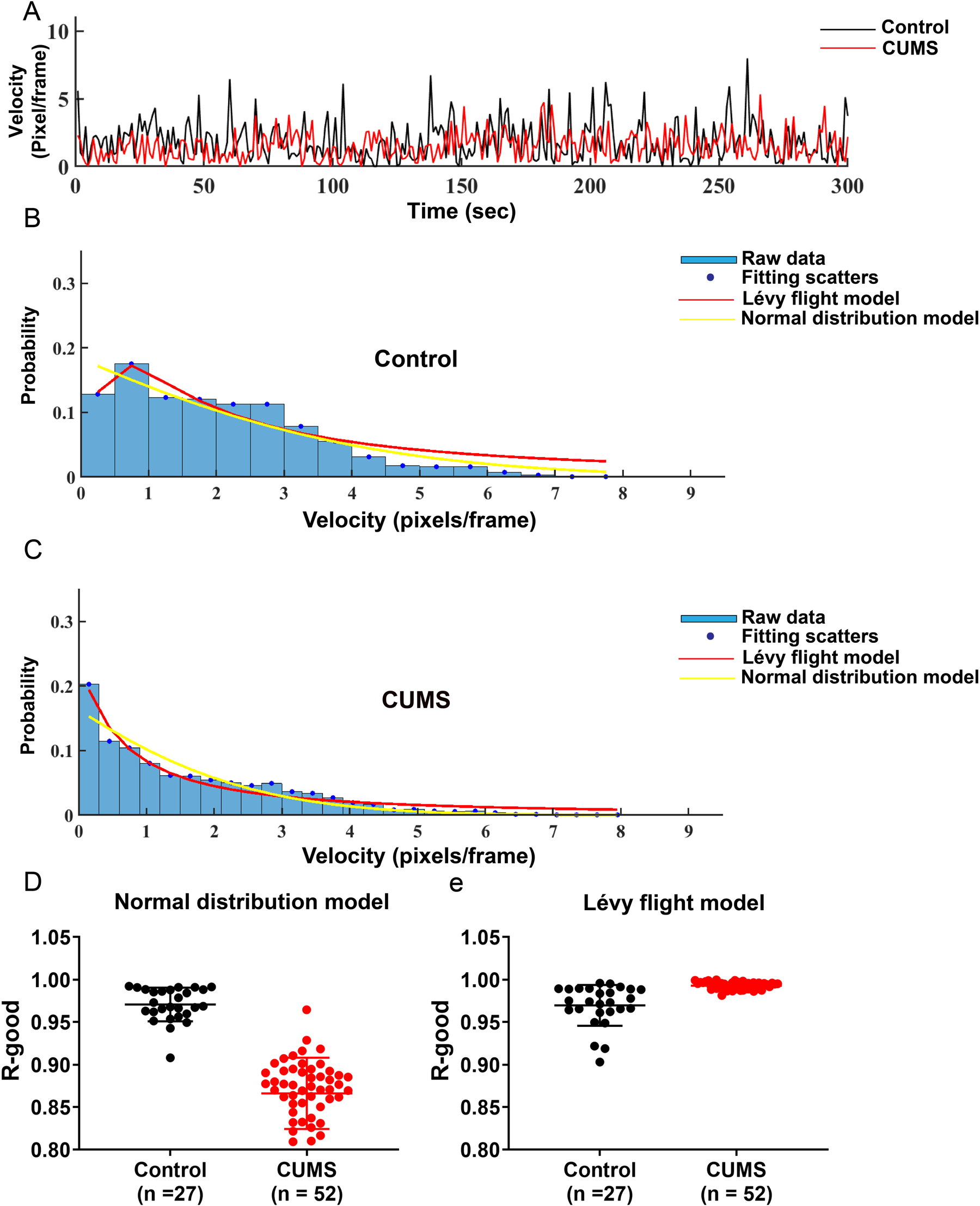
The walking velocity of mice exhibited a pattern of LF. (**A**) A mouse was selected randomly, and the walking speed of the two groups was recorded as a function of time. (**B-C**) Random selection was performed for a control mouse and a CUMS mouse to apply the normal distribution and LF distribution models. (**D-E**) The comparison between the r-good of the normal distribution (**D**) and LF distribution model (**E**) was performed in each group. The data were presented as means ± s.e.m., and the number of animals is indicated on the graphs.

### LF pattern in the OFT distinguished control and stress in mice

The significant difference between the control and CUMS groups led us to fit the LF distribution model to the walking speed of animals from each group at every experimental stage. A mouse’s walking speed distribution was represented by a curve with two parameters, γ, and μ (Equation 3, Method Section), and each mouse was denoted as a dot on the γ-μ plane. Control mice were represented with black dots, while CUMS mice on day 14 were depicted with red and blue dots in Figure 3A. The 2D Kolmogorov-Smirnov (2D K-S) test was utilized to determine whether the two-dimensional variables followed the same distribution. Results indicated a substantial difference between the control and CUMS groups (Figure 3A, *p* < 0.001). The CUMS-stressed mice were divided into CUMS + Saline and CUMS + FXT groups after day 14. On day 42, there was a significant difference between the CUMS + Saline and CUMS + FXT groups (Figure 3B, *p* < 0.001). Furthermore, compared to the control, the CUMS + Saline group exhibited more considerable differences during OFT (Figure 3B, *p* < 0.001). In contrast, no significant difference was observed between the control and CUMS + FXT group. Similar methods were used to compare OF data of the CUMS + Saline group on day 0, 14, and 42, revealing significant differences between day 0 and day 14 (Figure 3C, *p* < 0.001), day 14 and day 42 (Figure 3C, *p* = 0.040), and day 0 and day 42 (Figure 3C, *p* < 0.001). By comparing OF data differences at the three-time points in the CUMS + FXT group, significant differences were observed in pairwise comparisons at each time point (Figure 3D, *p* < 0.001). These results provide a quantitative approach to determine the mood state of CUMS mice.

**Figure 3.**
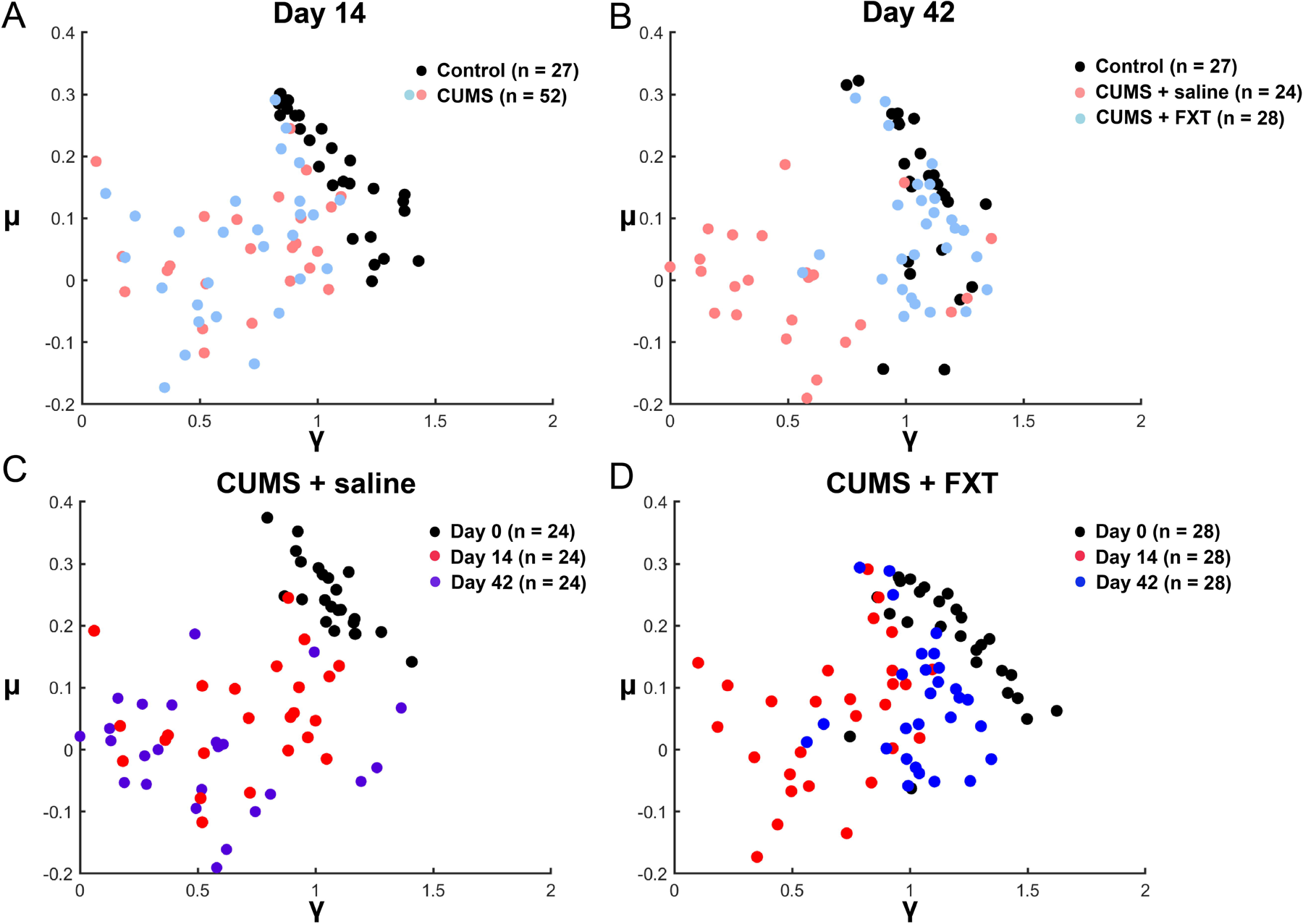
Each mouse was represented as a dot on the γ - μ plane. (**A**) The black dots on the γ-μ plane represented control mice (n = 27), while red and blue dots represented CUMS mice (n = 52). Our analysis indicated that LF patterns in the OFT distinguished Control and CUMS mice. The OF data for the two groups were analyzed at the 14th-day time point after CUMS modeling. A substantial difference between control and CUMS mice was observed on the γ-μ plane. (**B**) After day 14, CUMS-stressed mice were divided into CUMS + Saline and CUMS + FXT groups. On day 42, a significant difference between the CUMS + Saline and CUMS + FXT groups was observed. The comparison between Control and CUMS + Saline during OFT indicated a substantial difference, while no significant difference was found between Control and CUMS + FXT groups. (**C**) The OF data for the CUMS + Saline group were compared using the same method at 0, 14, and 42 days. d Significant differences were observed in pairwise comparisons of OF data at the three-time points in the CUMS + FXT group. The number of animals is indicated on the graphs. The null hypothesis that the 2D sets of variables followed the same distribution was tested using the two-dimensional Kolmogorov-Smirnov (2D K-S) test.

### Conventional and novel methods to differentiate between depression and anxiety in mice

The CUMS model can exhibit both anxiety-like and depression-like behaviors. In this study, the remaining mice underwent various classical neurobehavioral tests, including OFT, EPM, LDB, TST, and FST (Figure 1A). Following the methodology proposed by Shengyuan Yu et al. (see “Standard depression and anxiety disorder scales” section), we distinguished depression-like or anxiety-like behavior in mice. Initially, we normalized the depression/anxiety indicators of each experimental group to a scale ranging from 0 to 1. Subsequently, we assigned scores to each group based on the criteria provided in Table 1 Supplementary and Table 2 Supplementary. Finally, we comprehensively evaluated each mouse using the CRITIC method, with the weight ratios of the various behavioral indicators detailed in Figure 4A and Figure 4B. The overall CRITIC scores for each mouse are available in the online Experimental data section. Based on the scores, CUMS mice were categorized into four groups, with nine mice exhibiting anxiety-like behavior (Anxiety) (Figure 4C), seven mice showing depression-like behaviors (Depression) (Figure 4D), and eight mice with comorbid depression and anxiety (A & D). Upon re-examination of the behavioral indicators of the four groups following differentiation, it was observed that not all indicators could effectively differentiate between depression and/or anxiety (Figure supplement 2).

**Figure 4.**
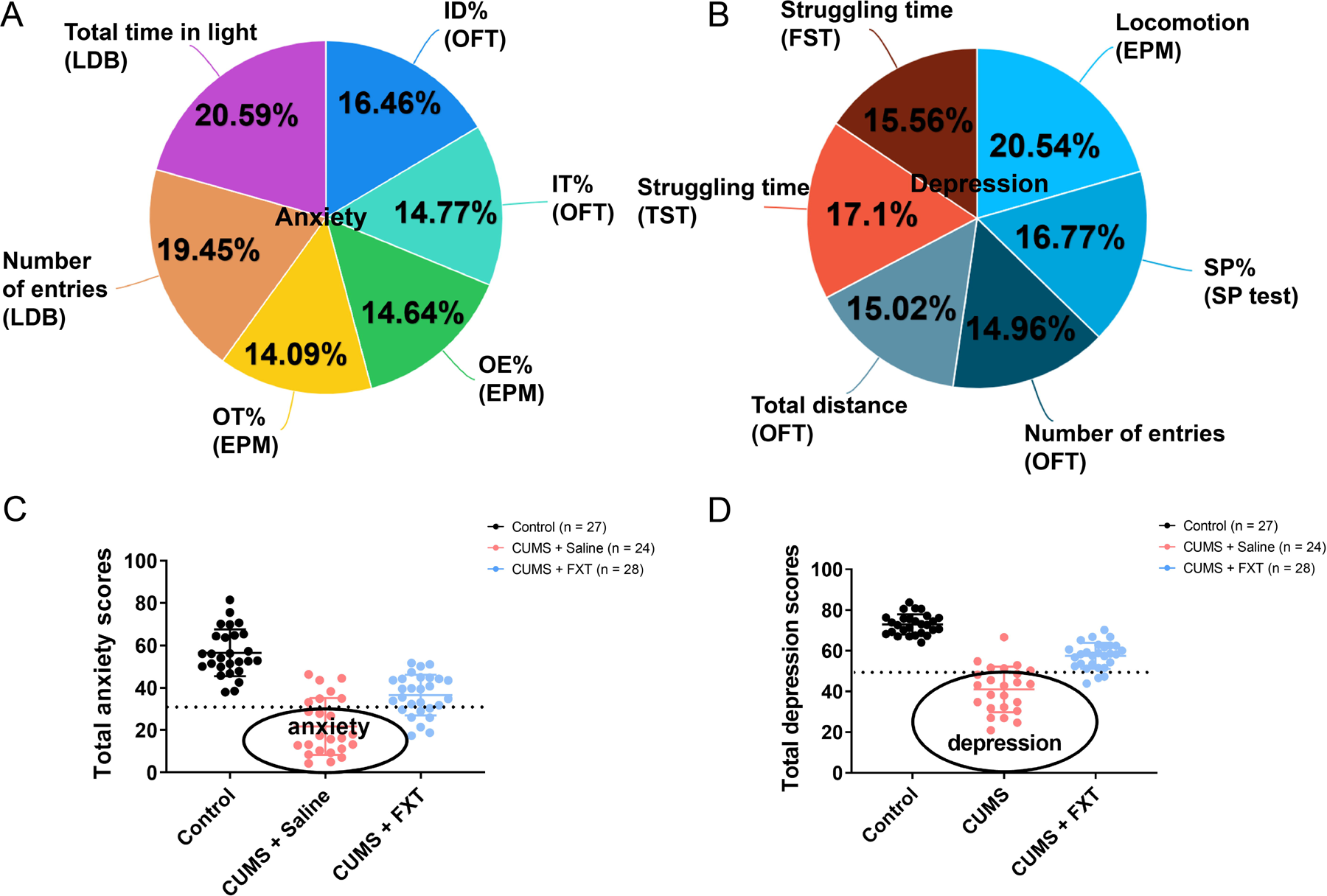
Mice exposed to CUMS exhibited typical symptoms of anxiety, depression, and comorbid anxiety and depression (A&D). (**A**) The proportion of anxiety indicators in mouse behavioral studies, as represented by the sector diagram, was determined using the criteria importance through the intercriteria correlation (CRITIC) method. (**B**) Using the CRITIC method, we calculated the sector diagram proportion of depression indicators in mouse behavioral studies. (**C**) Following the normalization of anxiety indicators within each group, the CRITIC-derived percentage was multiplied by the normalized value of each indicator and subsequently summed to obtain the total anxiety score for each mouse. Using the control group’s score as a reference point, mice with an anxiety score below 30 were deemed anxious in the CUMS group. (**D**) Upon normalization of depression indicators within each group, the CRITIC-calculated percentage was added to the normalized value of each indicator to obtain the total depression score for each mouse. Using the control group’s score as a reference, mice with a depression score below 30 were classified as depressed in the CUMS group. The data is presented as mean ± standard error of the mean (s.e.m); the number of animals is indicated on the graphs.

The LF fitting analysis was conducted twice on OFT video recordings, on the 14th and 42nd day of CUMS modeling, for the control, depression, anxiety, and A&D mice groups. On day 14, significant differences were observed between the control and the remaining three groups (*p* < 0.001, respectively). Notably, no significant difference was detected in the 2D K-S test between the Anxiety, Depression, and A & D groups (Figure 5A). On day 42, significant differences were observed between the control and the remaining three groups (Figure 5B, *p* < 0.001, respectively). Furthermore, significant differences were detected between Anxiety and Depression (Figure 5B, *p* = 0.001). The data for Depression and A & D also exhibited significant differences.

**Figure 5.**
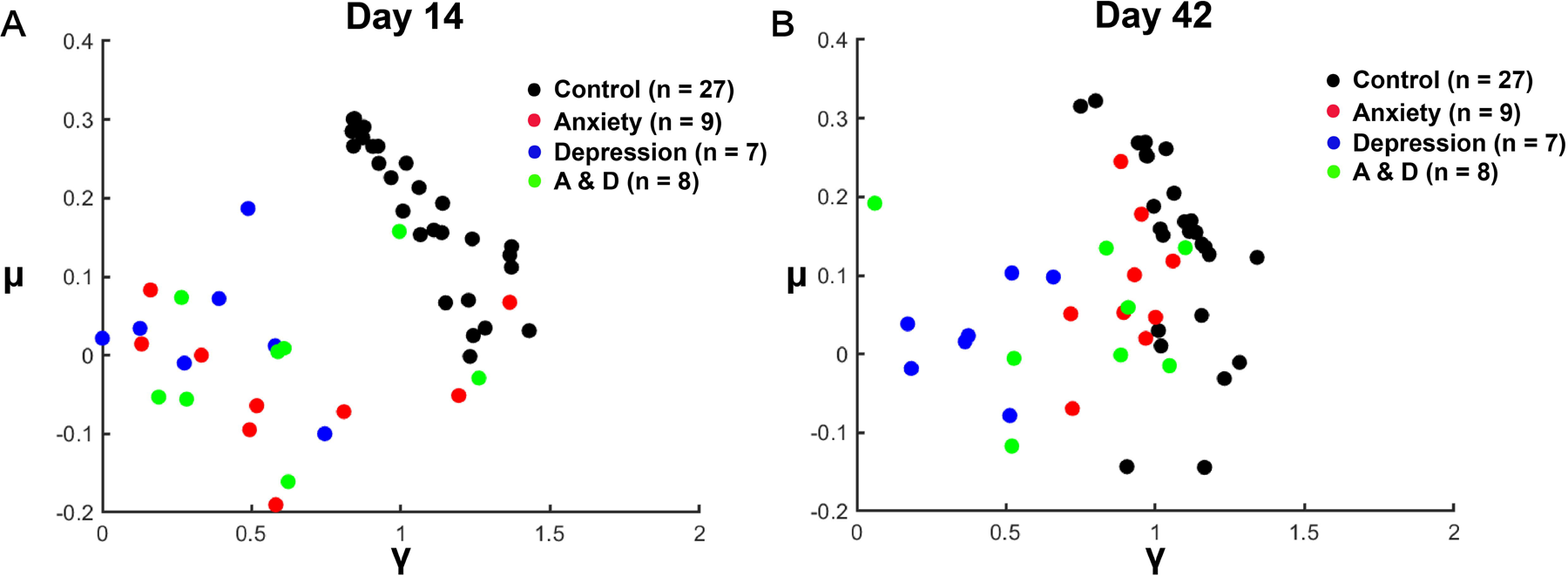
Assessment of the reliability of LF analysis in characterizing anxiety and depression in mice. (**A**) The black region indicates the data distribution range for the control mice. In contrast, the red, blue, and green regions represent the data distribution range for the anxiety, depression, and A&D mice, respectively, on day 14. Significant differences were observed between the control group and the other three groups (*p* < 0.001 for all comparisons) on day 14; however, no significant difference was observed in the two-dimensional Kolmogorov-Smirnov (2D K-S) test between the Anxiety, Depression, and A&D groups. (**B**) Significant differences were observed between the control and the other three groups on day 42 (*p* < 0.001 for all comparisons). Furthermore, significant differences were noted between the Anxiety and Depression groups (*p* = 0.001) and between the Depression and A&D groups (*p* = 0.009). The number of animals is indicated on the graphs. The two-dimensional Kolmogorov-Smirnov (2D K-S) test was employed to evaluate the null hypothesis that the 2D sets of variables conform to the same distribution.

### Preliminary LF sorting for anxiety and depression

Given the above results, we next focused on establishing a preliminary LF-based standard to sort mice into three groups, namely depression, anxiety, and healthy controls. We realized that multiple classical stress means existed to produce animal models. OF data obtained from our team’s previous experiments on animal models of depression and anxiety were collected for analysis. These models consisted of CUMS, ES (anxiety model), and CRS (depression model). A total of 99 mice exhibiting anxiety-like behavior were enlisted through the CUMS procedure, while 46 mice displaying depression-like behavior were enlisted using the same procedure. Additionally, 89 anxiety-like mice were enrolled through ES, and 69 depression-like mice were enlisted through CRS. A group of 237 control mice was also included in the study. Each diseased mouse underwent verification using traditional methods, and the LF procedure was performed on each mouse using the OFT. Their coordinates in the γ -μ plane were marked in Figure 5A. Given the data’s distribution, we used the support vector machine (SVM) algorithm to find the optimal separation of the three mice groups in the γ - μ planes. The optimal linear separation between the control and the anxiety was L_1_: μ = - 0.59γ + 0.79. The optimal linear separation between the depression and the anxiety was L_2_: μ = 6.54γ - 3.91 (Figure 6A).

**Figure 6.**
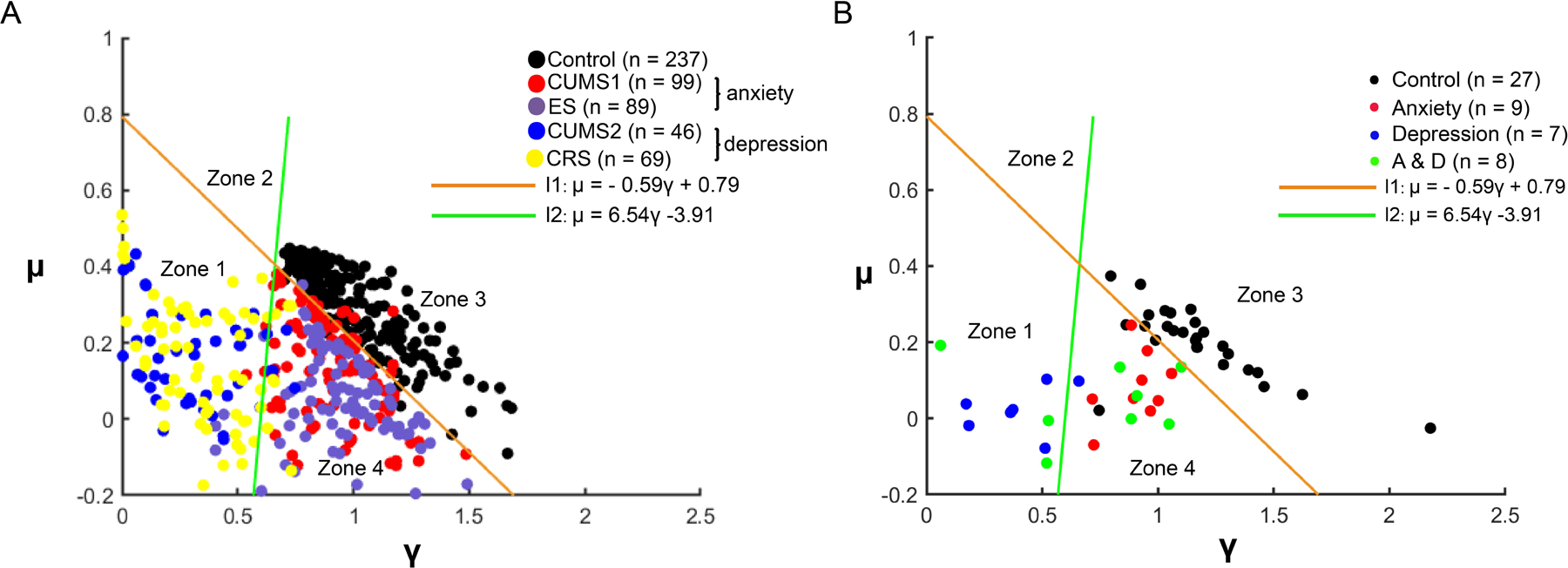
The LF method was employed to differentiate open field videos of mice exhibiting depression or anxiety, which were validated in prior experiments. (**A**) Three distinct models, namely chronic unpredictable mild stress (CUMS), electric shocks (ES), and chronic restraint stress (CRS) were utilized. CUMS was categorized into two groups based on behavior, anxiety-like (CUMS1; n = 99) and depression-like (CUMS2; n = 46). Additionally, ES (n = 89) represented an anxiety-like model, while CRS (n = 69) depicted a depression-like model. The control group (n = 237) was also included. Following successful modeling, LF fitting was performed on OFT videos to determine the precise values of γ and μ. A support vector machine (SVM) calculated two lines that distinguished the three data groups based on training data. The optimal linear separation between the control and anxiety-like groups was determined as L_1_: μ = - 0.59γ + 0.79, while L_2_: μ = 6.54γ - 3.91 represented the optimal linear separation between the control and anxiety-like groups. (**B**) The two-dimensional map distribution of the four groups of mice, validated in this experiment, was shown using two curves fitted by 540 mice. Rephrased sentence: The control group selected the data of OFT tested for the first time (Day 0), and the other groups selected the anxiety group, depression group and A&D group distinguished by CUMS modeling (Day 42). The number of animals is indicated on the graphs.

We used the L_1_ to distinguish between the control and the anxiety and the L_2_ to differentiate between the depression and the anxiety mice in this test. We found six mice in the depression group were in the depression-like behavior region, and all the anxiety mice were in the anxiety-like behavior region (Figure 6B).

## Discussion/Conclusion

The above results reveal a previously unknown statistical walking pattern of mice in anxiety and depression mental stages. Our findings revealed that mice exposed to CUMS displayed a characteristic LF velocity distribution. Furthermore, it was possible to distinguish mice exhibiting symptoms of anxiety or depression by analyzing the γ and μ parameters derived from LF statistics. This finding offers a novel, illustrative method to differentiate anxiety, depression, and healthy mice. This method is advantageous because it provides a one-step gentle approach (free walk in an OF) instead of the traditional multi-step stressful tests. This innovative approach has the potential to alleviate the industry’s long-standing concerns regarding the strong stress that traditional tests impose on animals. While some previous studies have also reported the behavioral features of stressed mice, few statistical models have been developed to help differentiate and quantify highlighted rodent models ^20^. For instance, Jin et al. reported that mice exhibiting anxiety-like behavior displayed a propensity for increased exploration of the outer area of the box while reducing their time spent in the central region ^21^. Belzung et al. mentioned that mice unable to escape in a confined space with increased immobility time have depressive-like behavior ^22^. These observations illuminate the correlation between rodents’ mental stages and their behaviors. However, when rodents exhibit mild symptoms or when the therapeutic benefits are not readily observable, identifying differences in behavioral markers associated with depression and anxiety can be a challenging task. Additionally, there are variations in the behavioral cues utilized to evaluate models of depression and anxiety ^23^. Of course, we should also consider that the order of tests may impact the experimental outcomes. To minimize additional stress on rodents, we plan to incorporate the OFT exclusively for assessing depression and anxiety after modeling. Additionally, we will employ the LF method to evaluate the stress levels of rodents. With our new approach, it is possible to process the OFT videos of rodents in an automated batch fashion, enabling the acquisition of quantitative results with minimal stress imposed on the animals.

When utterly ignorant of the orientation and distance of the prey, an animal may unintentionally choose a searching strategy among a few choices of styles. Ralf Metzler et al. calculated that searching trajectories that follow the LF statistics have higher efficiency than those following the normal statistics when the prey is relatively far from the initial starting location ^24^. On the contrary, if the prayer is initially close to the starting position, a normal statistics trajectory is of higher efficiency ^25^. Therefore, it is intuitive to assume that evolution has taught many animals a survival strategy such that a decision is made between the normal distribution and the LF distribution based on the expected prey distance. Animals in various stressed mental stages may have different prayer distance expectations and thus adopt various searching strategies. Our findings imply that the governing parameters of LF statistics may quantify a mental stage’s severity.

Lack of food is the most common stress animals face in nature. When food and other resources are abundant, mobile organisms tend to forage in a pattern of Brownian motion ^26^. In contrast, when faced with food and other resources shortage pressure, they often present an LF searching strategy, which means the distribution of traveling distances decays as a power law ^14,24^. The stress response of food deficiency in nature is similar to the fear of the unknown for the anxiety and depression mice. The anxiety and depression mice display heavy-tailed distributions in an open field and are consistent with the LF employed by animal predators searching for scarce food sources. In contrast, the speed of non-stress mice performed a normal distribution. A random walk is a nature-inspired algorithm consisting of consecutive unexpected steps or velocities. The LF distribution model usually exists in animal trajectories under survival pressure. This movement trajectory can effectively improve the survival rate when food is scarce. LF has also been applied to other research fields related to bionic ^25,27^. Although little is known about how LF searching systems developed, evidence indicates that this strategy may have evolved to allow animals to cover the most expansive area with the least energy, which may explain the widespread occurrence of such patterns among modern animals. We suspect that this pattern of behavior, imprinted in animal brains by survival advantage in evolution, should also exist in laboratory animals.

FXT has been well-reported to alleviate the symptoms of CUMS ^28^. As anticipated, it was observed that the administration of the drug caused a shift in the mice’s behaviors on the γ - μ plane toward the healthy region from Figure 3A towards Figure 3B (blue dots). On the other hand, the symptoms of the CUMS + saline group appear to be more severe (as indicated by the pink dots). Next, we conducted a time-based analysis of the behavioral tests of each mouse in the group identified as the CUMS + saline group. Figure 3C illustrates that their positions on the γ – μ plane shifted gradually from the upper right region (black) to the middle region (purple) and then to the lower left region (red) through time, indicative of an increase of severity in their symptoms. Similarly, the behaviors of the CUMS + FXT group were analyzed over time, and it was observed that they initially resided in the upper right region (black) of the γ – μ plane, which then transitioned to the middle region (red), and eventually recovered to the upper right region (blue). These observations suggest that our quantification method is capable of demonstrating the effects caused by positive drug administration. With further rigorous validations in the future, there is reason to be optimistic that the length of the dot-shifting vector in the γ-μ plane could serve as a quantitative measure of a drug’s therapeutic efficacy.

In this work, we proposed for the first time that LF can distinguish depression and anxiety mice in an OF test. Furthermore, we used speed as a surrogate indicator since stressed animals should subconsciously increase their speed after activating LF search patterns. Finally, we found that the velocity distribution of the stressed mice in the OF matched that of LF. The r-good value was significantly different from that of the normal mice. At the same time, depression and anxiety groups of mice also showed significant differences in γ and μ parameters, which we could easily distinguish^29^. Based on the velocity LF fitting pattern of the stressed mice in OF, we could qualitatively evaluate the anxious or depressive state of the tested mice and the efficacy of novel therapeutic agents in a rapid, straightforward, and low-cost way.

In the upcoming discussion, we aim to analyze the significance of the parameters associated with the LF distribution. Given the inherent abstractness of these parameters, they may be challenging to visualize, thereby warranting a need for appropriate visualization methods. To this end, we propose using a visualization technique known as “typical mice” to represent each category and demonstrate the respective groups’ statistical behavior. To construct the typical healthy mouse, we computed the average γ and μ parameters of the control mice group, yielding γ = 1.182 and μ = 0.205, respectively. This typical mouse was named Healthy Doe. Similarly, we derived the Depress Doe (γ = 0.380 and μ = 0.005) and the Anxiety Doe (γ = 0.917 and μ= 0.095) by computing the average values of the corresponding groups. These parameters are plugged into Equation 3 in the Method section and the typical velocity distributions of each group are plotted. In order to plot the typical velocity distributions for each group, the γ and μ parameters obtained for the respective group were inputted into Equation 3, as described in the Method section. The resulting output from this equation provides a graphical representation of the distribution of velocities exhibited by each group. These distributions were then plotted and can be used to compare and contrast the statistical behavior of each group (Figure supplement 3).

A hitherto undiscovered statistical gait pattern in mice exhibiting symptoms of anxiety and depression is unveiled by this investigation. These results introduce a novel and uncomplicated framework for distinguishing between anxiety, depression, and healthy mice. Contrary to conventional and burdensome multi-step tests, this method offers a non-invasive single-step approach (unrestrained ambulation in an open field) that is gentle in nature.

## Materials and Methods

### Animals

Experiments were conducted using 145 male and female C57Bl/6 mice (6-8 weeks old, weighing 18-24 g) obtained from Charles River Laboratories Animal Technology Co., Ltd. (Beijing, China). The animals were housed under specific pathogen-free conditions, provided standard laboratory chow and distilled water, and maintained on a 12-hour light-dark cycle in standard cages with corn cobs in a room maintained at an ambient temperature of 23 ± 1 ℃ at Kangcheng Biotech, Ltd, Co. (Sichuan, China) facilities [animal production license number: SYXK (Chuan) 2019-215]. All procedures followed the guidelines of the Association for Assessment and Accreditation of Laboratory Animal Care (AAALAC) and were approved by the Institutional Animal Care and Use Committee (IACUC) of the West China Hospital, Sichuan University (Approval No. 2019194A).

### A mice model of chronic unpredictable mild stress-induced depression or anxiety

The chronic unpredictable mild stress (CUMS) modeling process was adjusted slightly. It consisted of sequential application of mild stressors, including cage tilting, heat stress, noise stimulation, wet bedding, loneliness, fasting, inversion of a day-night cycle for 24 hours, water deprivation, reciprocating sway, random foot shock for 120 seconds, crowding, cold stress, flash stimulation for 12 hours, constraint for 4 hours, and cage replacement for 24 hours. Each stimulus was randomly arranged, and two different stressors were performed consecutively daily throughout the experiment ^6^.

### A mice model of chronic restraint stress-induced depression

The chronic restraint stress (CRS) protocol underwent slight modifications compared to previous descriptions^30,31^. The establishment of a mouse model of depression often involves the common utilization of CRS. One week before the start of stress exposure, animals were divided into control and experimental groups to acclimate to new cages. Mice were placed in a cylinder for 4 hours daily (8:00-12:00) for 14 consecutive days, almost immobilizing them. The size of the cylinder matched the size of the animal. Non-stressed controls were moved to a test room and handled gently for 5 minutes before returning to their holding room 4 hours later.

### A mice model of electric shocks-induced anxiety

The electric shocks (ES) modeling session was executed following prior descriptions, incorporating minor modifications. Electric foot shocks were employed to induce an anxiety mouse model ^32^. After a 30-minute adaptation, mice underwent ten intermittent inescapable electric foot shocks delivered by an isolated shock generator (Shanghai Jilang Information Technology Co., Ltd, China) through the grid floor. The shocks had an intensity of 0.8 mA, an interval of 15 seconds, and a duration of 15 seconds. Control mice were placed in the same chamber for 5 minutes without undergoing electric foot shock.

### Grouping

Before CUMS induction, all mice underwent a sucrose preference (SP) test and an open field test (OFT) test to preclude physically and mentally low activity subjects. Before the experiment (day -2), we evaluated the sucrose preference of mice. Those with a preference rate lower than 80% were excluded, removing 11 mice. At the onset of the modeling process, the CUMS modeling group was assigned the designation D0. On Day 0, the remaining 134 mice underwent an open field test, excluding 31 that made fewer than five entries into the center zone. Subsequently, a total of 103 mice were allocated into two groups, namely the control group comprising 32 mice and the CUMS modeling group consisting of 71 mice (as the CUMS modeling group will later be divided into CUMS + Saline group and CUMS + Fluoxetine group). On D14, we conducted model validation by performing additional tests on SP and the OFT. Control mice with a sucrose preference percentage lower than 80% and CUMS mice with a ratio higher than 80% were excluded from the SP test. In the OFT, CUMS mice that crossed the center zone more than five times were also excluded. Unfortunately, throughout the experiment, we experienced two deaths in the control group and four deaths in the CUMS group. Consequently, we had a total of 27 remaining mice in the control group and 52 successfully modeled mice in the CUMS group. On D14, as part of the model validation process, we performed additional sucrose preference (SP) and open field test (OFT) experiments. Control mice with a sucrose preference percentage below 80% and CUMS mice with a sucrose preference percentage above 80% were excluded from the SP test. In the OFT, CUMS mice that crossed the center zone more than five times were excluded. Regrettably, during the experiment, two deaths occurred in the control group, and four deaths occurred in the CUMS group. Consequently, there were a total of 27 mice remaining in the control group and 52 successfully modeled mice in the CUMS group. Following Day 14, we partitioned the CUMS group into CUMS + Saline and CUMS + Fluoxetine (FXT) cohorts. The CUMS + Saline group was administered 5mL/kg of Saline each day after modeling. In contrast, the CUMS + FXT group was given a daily dosage of 20mg/kg of Fluoxetine hydrochloride after modeling. Treatment was sustained until the culmination of the experiment (The pertinent data is available within the supplementary materials, Figure supplement 1). To ascertain the range of Control, Anxiety, and Depression mice on the γ-μ plane; we utilized the pre-established CUMS, chronic restraint stress (CRS), and electric shocks (ES) models in our laboratory. These models effectively discriminated between mice displaying anxiety-like behavior and those exhibiting depression-like behavior (see method: Standard depression and anxiety disorder scales). The OFT data encompassed 540 mice in the control and model groups, with supplementary information on the 540 mice data available in the online supplementary materials.

### Physical health assessment

Body weight, food intake, and coat state score were evaluated weekly, specifically on Wednesdays between 10 am and 12 am, until the conclusion of day 49. The 24-hour food consumption was meticulously documented. The assessment of the coat state score encompassed seven distinct areas, namely the head, neck, dorsal region, ventral coat, tail, anterior claw, and hind claw. This evaluation aimed to identify any signs of deterioration in the physical condition of the coat, such as fur loss and accumulation of dirt^33^.

### Sucrose preference (SP) test

In preparation for the SP test, 1% sucrose solution and distilled water were placed simultaneously in each mouse’s cage two days before the test for taste adaptation. On the third day, following a six-hour fast and water deprivation, mice were raised individually in cages with one pre-weighed bottle of 1% sucrose solution and another of distilled water. After six hours, the positions of the two bottles were interchanged. Subsequently, after an extra twelve hours, both bottles were eliminated, and the residual liquid was quantified to ascertain mouse sucrose preference, employing the subsequent Equation 1 ^34^.

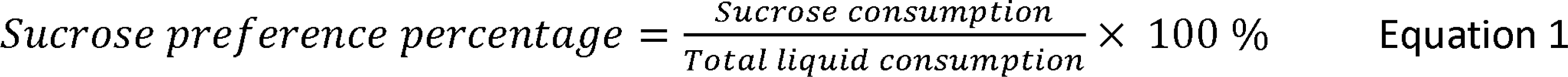

### Behavioral assessment

The mice were acclimated to the testing room for 12 hours before assessment. Maintaining a clean, odor-free, and quiet environment was crucial for the experiment, and the operator remained concealed during testing. The Topscan Package (Clever Sys Inc., USA) was used to record and analyze mouse movements.

### Open field test (OFT)

Each mouse was gently placed in a 50 * 50 * 50 cm OF box for 5 minutes to explore freely under weak (50 lx) illumination. The total distance traveled (cm), and number of entries into the central area (25 * 25 cm) were recorded and analyzed using the software. IT% was calculated as the time spent in the inner area/300 s * 100, while ID% was calculated as the inner area distance/total distance * 100 ^35^.

### Elevated plus-maze test (EPM)

The EPM had two open and two closed arms. Each arm measured 30 cm in length and 5 cm in width, with walls that were 20 cm high in the closed arms. The EPM was elevated to a height of 60 cm above the ground, with a 90°angle between the arms. Prior to the test, mice were placed in the central area facing the open arm and allowed to explore for 5 minutes. Open Arm Entries Percent (OE%) was calculated as Entries into the Open Arm divided by (Entries into the Open Arm + Entries into the Closed Arm) multiplied by 100. Open Arm Time Percent (OT%) was calculated as time in the Open Arm divided by 300 seconds multiplied by 100 ^36^.

### Light/dark box test (LDB)

Anxiety was tested in the LDB as previously described. Each mouse was placed in an apparatus consisting of two identical opaque Plexiglas compartments (18 * 12 * 12 cm) and an opening (5 cm * 5 cm). The mice were positioned in the light box with their heads facing the dark box, and their exploratory behavior was monitored for 10 minutes following their initial crossing ^37^.

### Tail suspension test (TST)

The mice were suspended by their tails using adhesive tape hooked onto a horizontal rod. The distance between the tip of the mouse’s nose and the floor was approximately 25 cm. After being suspended for 6 minutes, the time spent immobile during the last 4 minutes was recorded ^12^.

### Forced swimming test (FST)

Before testing, the mice were placed in plastic cylinders filled with water (25 ± 1 ℃; depth: 15 cm) for 15 minutes. The next day, the mice were placed in a 10-cm-wide cylinder (height: 30 cm), and the time spent immobile during the final 4 minutes of the 6-minute forced swimming test was recorded^38^.

### Fluoxetine

Fluoxetine hydrochloride (FXT, Sigma Chemical Company, St. Louis, MO, USA) was dissolved in Saline and administered once daily via the intraperitoneal route for 37 days at 20 mg/kg. The fluoxetine hydrochloride dosage was selected based on previous studies ^39^.

### Standard depression and anxiety disorder scales

The depression and anxiety disorder scales in the rodent model were subtly modified by prior descriptions ^23^. In the first step, the outcome measures for each test were normalized to acquire a “measure score” ranging from 0 (indicating low anxiety/depression) to 1 (indicating highly anxious/depression) for each individual, as calculated using Equation 2 ^40^:

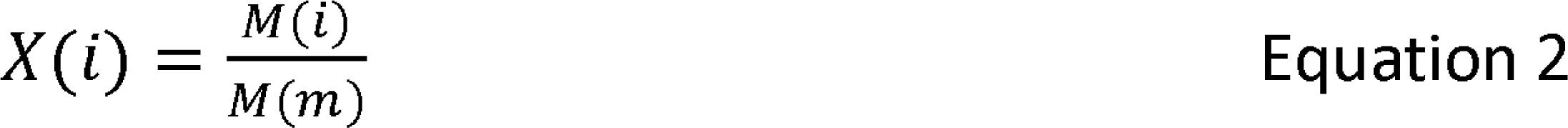

In Equation 2, the normalized individual measure score, denoted as X(i), is calculated based on the actual individual measure datum (M(i)), such as the number of entries, and the maximum measure datum (M(m)) observed in the study cohort (all mice).

The anxiety behavioral indices (ID%, IT%, OT%, OE%, time spent in the light box, number of entries) and the depression behavioral indices (SP%, number of entries in the central area, total distance, locomotion in closed arms, struggling time of TST, struggling time of FST) were subjected to the normalization process within each group of mice. The normalized scores were obtained from the behavioral indicators in Tables 1 and 2 in the Supplementary Material. To comprehensively assess each indicator of the mice, we employed the Criteria Importance Through Inter-criteria Correlation (CRITIC) method. This method was utilized to determine the objective criteria weights based on the experimental data, eliminating any influence from decision-makers and ensuring optimal conditions for both the strength and width ^41^. The specific comprehensive scores for each mouse, derived from the CRITIC method using anxiety and depression indicators, can be found in the online Experimental data. Based on the CRITIC composite scale, mice scoring below 30 were classified as anxiety mice, while those scoring below 50 were classified as depressed in comparison to the control group mice.

### LF fitting and Lévy distribution

The mice were placed in a standard OF apparatus for behavioral studies (Clever Sys., Inc. US), and their behavior was recorded through a series of videos. The recorded videos were analyzed using a custom MATLAB script (Math Works Inc. US, version R2019a). The mice were segmented from the videos frame-by-frame, and the mean coordinates of the segmented pixels were found to determine the center of each mouse. Walking speed was calculated by measuring the distance a mouse traveled between a specific number of frames, divided by the time interval. To maintain balance between measurement accuracy and sensitivity to speed changes, speed was calculated every five frames; we fitted two probability density functions (PDF) for the walking speed statistics of each mouse from all the categories. The first PDF is the LF distribution. A simplified and often employed form of the LF distribution is defined as ^26^:

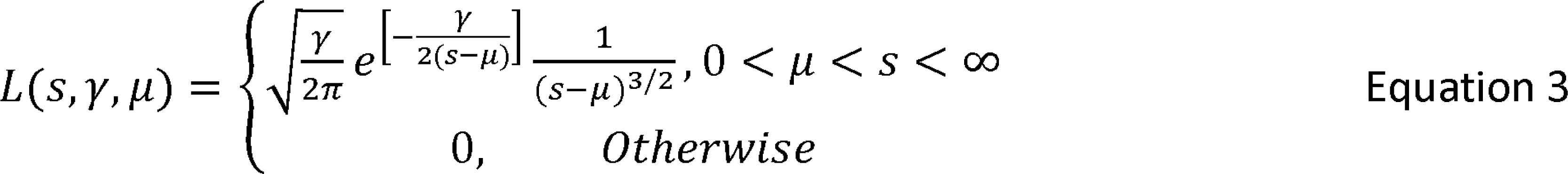

L is the probability density, µ > 0 is the minimum possible speed, and γ is a scale parameter.

We perform a parametric curve fitting of the distribution data to Equation 3. This method enables us to convert an animal’s random walk into a pair of parameters; hence, it can be plotted as a dot in a two-parameter γ-μ plane.

The other PDF is the normal distribution ^26^:

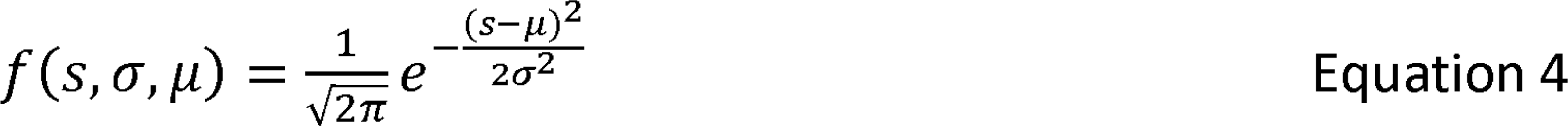

Statistical distributions can exhibit diverse shapes, yet they frequently adhere to typical patterns. Among these patterns, the normal distribution is particularly prevalent, characterized by a majority of values clustering around the mean and fewer values as they deviate from it, often referred to as the “tail” of the distribution. Both models depict random walks with distinct probability distributions in the context of LF and the normal distribution. Notably, the LF model possesses a distinctive feature whereby its distribution values in the far right region are slightly larger than those of the normal distribution.

The Goodness of Fit (R-good) is a quantitative representation of the curve fitting quality, which is defined as:

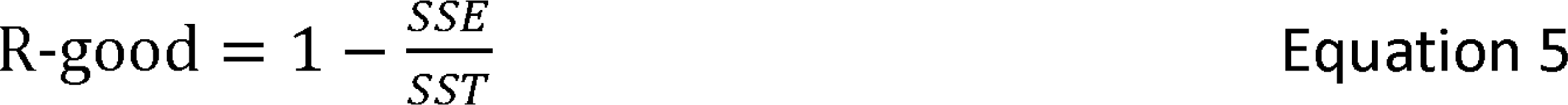

Where SSE (error sum of squares) is defined as 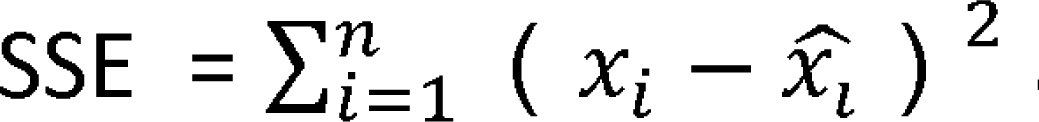 and SST is defined as 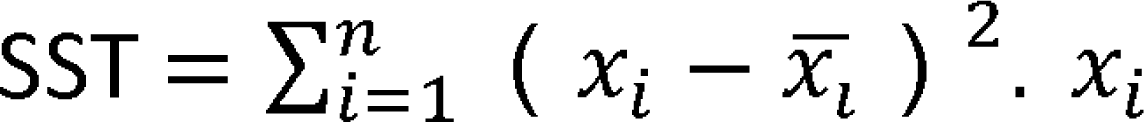 is defined as the average value of this set of data. The value of R-good is between 0 and 1. The closer it is to one, the better the fit is.

### Support vector machine (SVM)

The SVM is an algorithm used for classification. Its main goal is to find the best possible boundary that separates different clusters of data points in a way that maximizes the margin between them. It is suitable for the feature reduction of high-dimensional data with a small sample size. A general mathematical model of support vector machines is shown in Equation 6 and Equation 7.

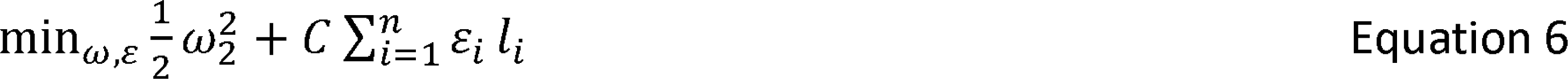

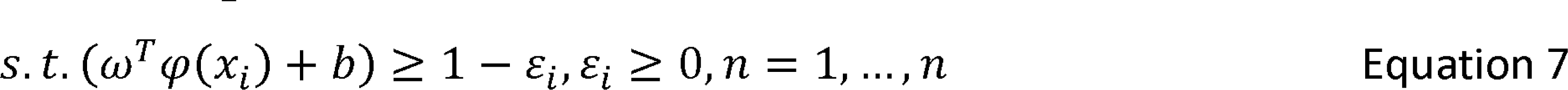

The detailed solution process of the SVM problem is found in Ref ^42^.

### Statistical information

The statistical analysis was done using IBM SPSS Statistics version 26.0 software (IBM Ltd., United Kingdom). Data that followed a normal distribution were presented as mean ± standard error of the mean (SEM). Unpaired t-tests were employed to compare the differences between the control and model groups. When the data followed a normal distribution and had equal variances, a one-way Analysis of Variance (ANOVA) followed by post hoc Tukey Dunnett’s multiple comparison tests was used to analyze differences among the model group, buspirone group, and each treatment group. A repeated measures ANOVA was conducted to examine variations in body weight, food intake, coat state, sucrose preference, and fecal amount among groups. The interaction effects between repeated indicators and days were assessed using Pillai’s trace. If interactions were observed between a specific variable and days, the differences among each group were compared at the final time point. If no interactions were present, post-hoc analysis using Bonferroni’s multiple comparison tests was performed. The correlations between each effect were evaluated using Pearson’s correlation. Statistical significance was set at a two-sided p-value < 0.05.

The 2-Dimensional Kolmogorov–Smirnov (2D K-S) test is a nonparametric, bivariate statistical test designed to verify whether two two-dimensional samples follow the same distribution ^43^. The point defined four quadrants for any given point in a two-dimensional space. The integration probability is defined as the probability of sample data distribution in one of the quadrants that accounted for the total number of samples. By iterating over all data points and quadrants, the test statistic D_FF,1_ is defined by the maximal difference of the integrated probabilities between samples in any quadrant for any origin from the first sample. Similarly, after going through all points of another sample, the test statistic D_FF,2_ is obtained. D_FF,1_ and D_FF,2_ are then averaged to compute the overall D_FF_ for hypothesis testing, D_FF_=(D_FF,1_+D_FF,2_)/2.

In the large sample limit (n ≥ 80), it was shown that D_FF_ converged in distribution ^43^.

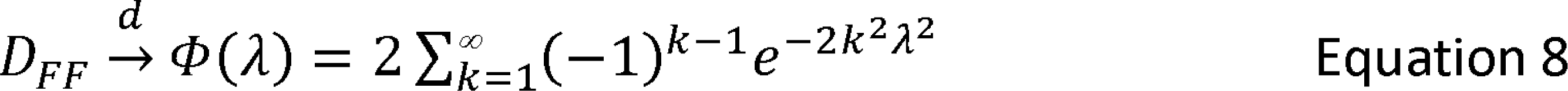

For a single sample of size n

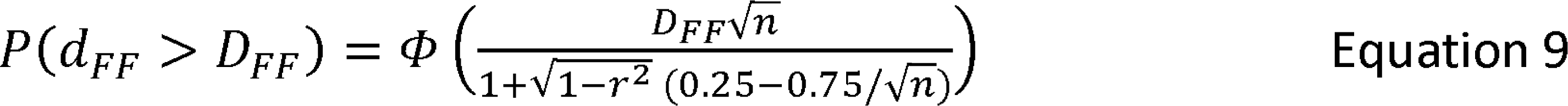

The two-sample case used the same formula as above, where n is defined as

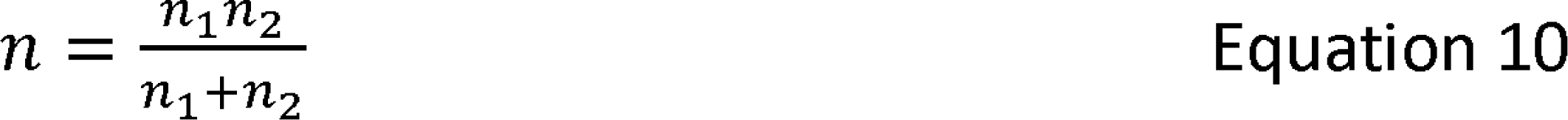

and r is defined as

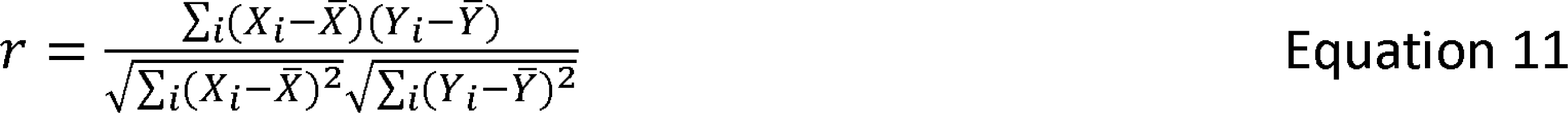

### Statement of Ethics

The Association for Assessment and Accreditation of Laboratory Animal Care (AAALAC) reviewed and approved this study protocol. It was approved by the Institutional Animal Care and Use Committee (IACUC) of the West China Hospital, Sichuan University (Approval No. 2019194A).

## Acknowledgements

This study was supported by the National Natural Scientific Foundation of China (82071349, 82027808, and 81771310), the West China Hospital of Sichuan University Discipline Excellence Development 1・3・5 Engineering Project (Interdisciplinary Innovation Project), the National KEY RESEARCH and Development Program (2017YFA0505903), and the National Science and Technology Ministry of China (2022YFF0706500 and 2019YFE0196700).

## Author Contributions

Q.L. coordinated the experimental operations and data analysis and assisted in preparing the manuscript. Y.N. conducted all statistical analyses and wrote computer code for this study. Y.L. and B.Z. handled the processing of all video data. N.Z. and W.K. provided practical psychiatric interpretations for the study. C.T. carried out the CUMS, ES, and CRS experiments. D.C. provided valuable input to the manuscript. Y.Z. helped to design the neurobehavioral experiments and supervised the data analysis.Z.W. co-designed the entire survey with Z.Z. and directed the mathematical research. Z.Z. conceived the original idea, designed, and supervised the entire study. All authors read and corrected the article.

## Data Availability Statement

The data can be available at the editor’s request, and experimental data has been uploaded separately. Further inquiries can be directed to the corresponding author. The data that support the findings of this study are not publicly available due to privacy reasons but are available from the corresponding author upon reasonable request.

