## Supplementary figure legends for "Lévy statistics define anxiety and depression in mice subjected to chronic stress"

### **Supplementary materials**

Supplementary figure legends

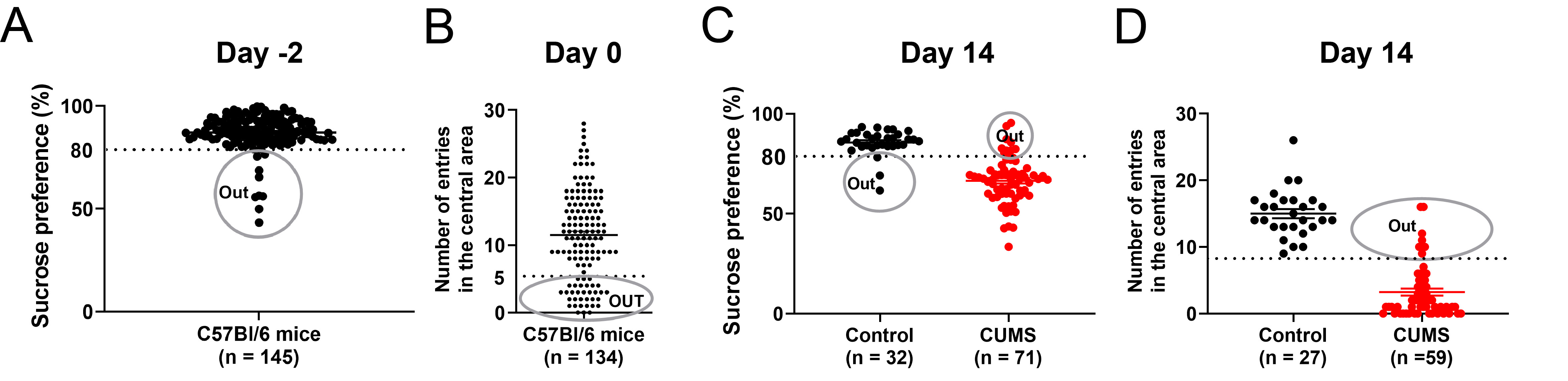

**Figure 1—figure supplement 1.** The CUMS data was collected both before and after the modeling process.

(**A**) To assess anhedonia in mice, the sucrose preference (SP) was evaluated prior to the experiment on Day -2. Mice with a sucrose preference rate of less than 80% were excluded from the experiment, and a total of 11 mice were eliminated. (**B**) On Day 0, physical evaluations of mice were conducted using the OFT. Mice with fewer than five entries in the center were excluded from the experiment, leading to the elimination of the thirty-one mice. (**C**) To assess anhedonia in mice, the SP was evaluated following CUMS. Mice with a preference rate of less than 80% were excluded from the CUMS group, resulting in the elimination of eight mice. However, the control group only had three mice eliminated. (**D**) On Day 14, the CUMS mice with fewer than five entries in the center in the OFT were excluded, leading to the elimination of seven mice. Regrettably, six mice, comprising two from the control group and four from the CUMS group, succumbed to starvation or mortality during the course of the experiment. Therefore, the final number of animals included 27 in the control group and 52 in the CUMS group. The CUMS mice exhibited a shorter travel distance than the unstressed controls (F_26,51_ = 1.389, *p* < 0.001 for Control vs. CUMS). The presented data represent mean ± s.e.m. Statistical significance was established when p values were below 0.05. The number of animals is indicated on graphs. Two-group comparisons were evaluated with unpaired t-tests.

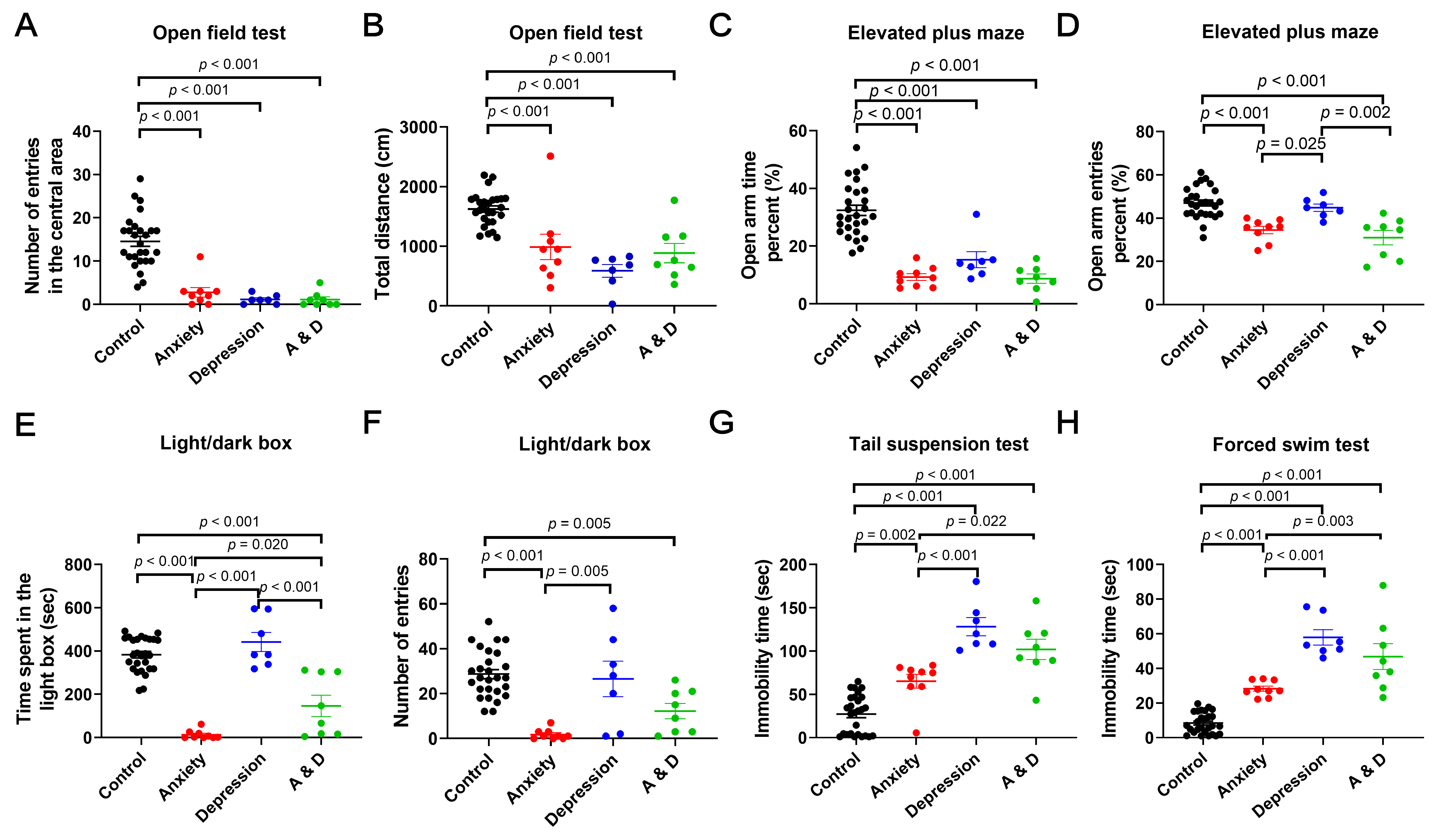

**Figure 2—figure supplement 2.** Behavioral phenotypes were assessed after grouping.

(**A-B**) On day 42, the Anxiety, Depression, and A&D groups displayed reduced central region crossings and traveled similar distances compared to the control groups (*p* < 0.001 for all comparisons).(**C**) In the EPM test, the Control mice exhibited significantly greater exploration time within the open arm relative to the other groups (*p* < 0.001 for all comparisons).

(**D**) Compared with the Depression group, the Anxiety and A & D groups entered the open ratio fewer times ( *p* = 0.025 for Anxiety vs. Depression, *p* = 0.002 for A & D vs. Depression). (**E-F**) In the LDB test, the Anxiety mice spent significantly lower time and less entries in the light box (*p* < 0.001 for Control vs. Anxiety). (**G-H**) In the TST and FST, the Depression and A & D mice showed the much more time for immobility time (*p* < 0.001, respectively). The data is presented as mean ± standard error of the mean (s.e.m); statistical significance was considered with p values less than 0.05. In the present study, the population of animals examined was composed of Control (n = 27), Anxiety (n = 9), Depression (n = 7), and A & D (n = 8) groups. Differences among the groups were evaluated using one-way ANOVA, followed by post hoc Bonferroni analysis.

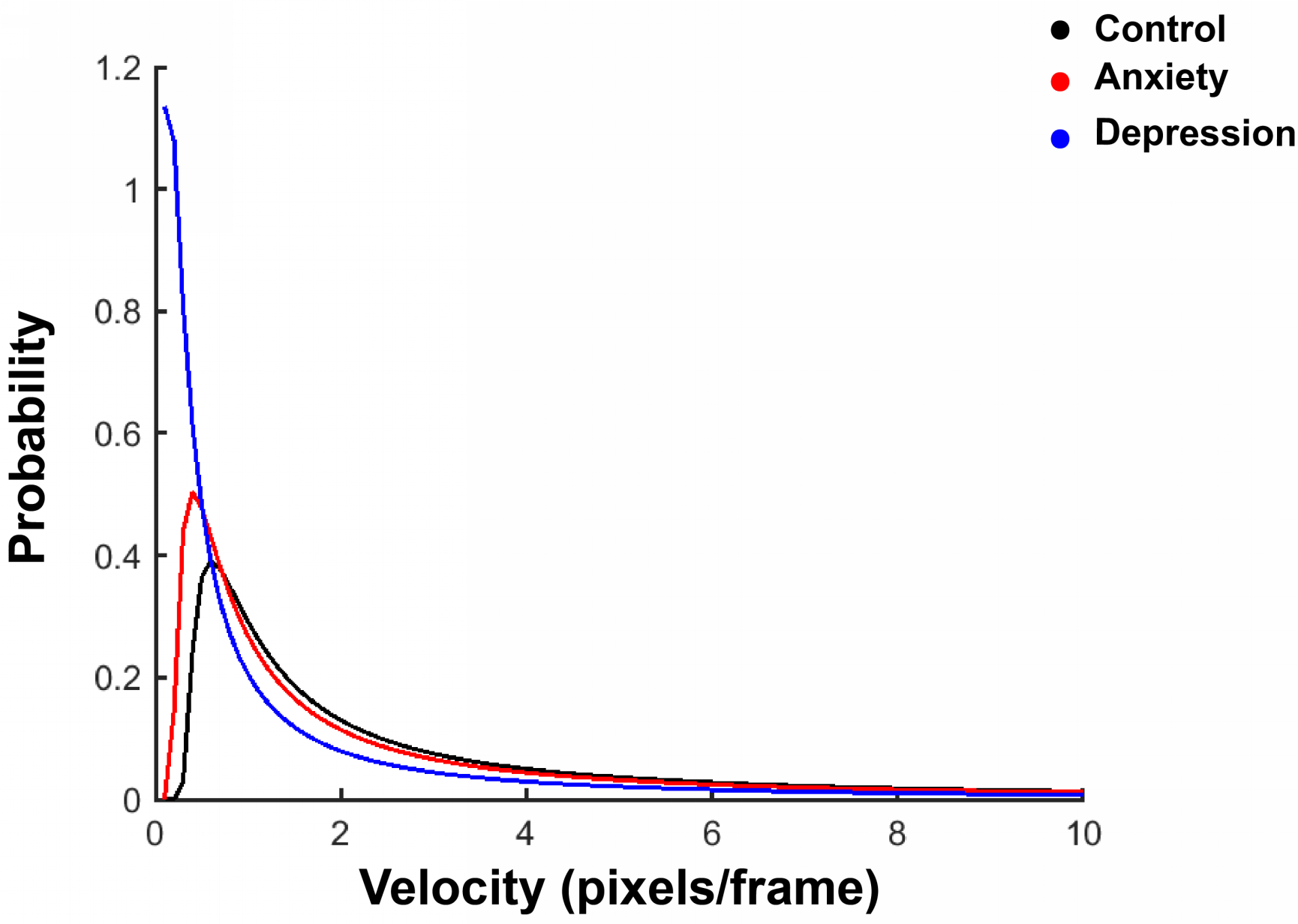

**Figure 3—figure supplement 3.** Probabilistic and velocity parameters

Probabilistic and velocity parameters were obtained by averaging miu for each Lévy flight (LF) model and averaging gamma. The resulting averages were then entered into the LF function to generate a set of curves. Three sets of three curves were drawn in this manner.

**Supplementary Table 1. Assessing the level of anxiety in mice.**

| **Anxiety indexes** | **Interval** | **Scores** |
| --- | --- | --- |
| **Inner zone distance percent, ID (%)** | 0-0.1 | 10 |
|  | 0.1-0.2 | 20 |
|  | 0.2-0.3 | 30 |
|  | 0.3-0.4 | 40 |
|  | 0.4-0.5 | 50 |
|  | 0.5-0.6 | 60 |
|  | 0.6-0.7 | 70 |
|  | 0.7-0.8 | 80 |
|  | 0.8-0.9 | 90 |
|  | 0.9-1 | 100 |
| **Inner zone time percent, IT (%)** | 0-0.1 | 10 |
|  | 0.1-0.2 | 20 |
|  | 0.2-0.3 | 30 |
|  | 0.3-0.4 | 40 |
|  | 0.4-0.5 | 50 |
|  | 0.5-0.6 | 60 |
|  | 0.6-0.7 | 70 |
|  | 0.7-0.8 | 80 |
|  | 0.8-0.9 | 90 |
|  | 0.9-1 | 100 |
| **Open arm time percent, OT (%)** | 0-0.1 | 10 |
|  | 0.1-0.2 | 20 |
|  | 0.2-0.3 | 30 |
|  | 0.3-0.4 | 40 |
|  | 0.4-0.5 | 50 |
|  | 0.5-0.6 | 60 |
|  | 0.6-0.7 | 70 |
|  | 0.7-0.8 | 80 |
|  | 0.8-0.9 | 90 |
|  | 0.9-1 | 100 |
| **Open arm entries percent, OE (%)** | 0-0.1 | 10 |
|  | 0.1-0.2 | 20 |
|  | 0.2-0.3 | 30 |
|  | 0.3-0.4 | 40 |
|  | 0.4-0.5 | 50 |
|  | 0.5-0.6 | 60 |
|  | 0.6-0.7 | 70 |
|  | 0.7-0.8 | 80 |
|  | 0.8-0.9 | 90 |
|  | 0.9-1 | 100 |
| **Time spent in the light box (sec)** | 0-0.1 | 10 |
|  | 0.1-0.2 | 20 |
|  | 0.2-0.3 | 30 |
|  | 0.3-0.4 | 40 |
|  | 0.4-0.5 | 50 |
|  | 0.5-0.6 | 60 |
|  | 0.6-0.7 | 70 |
|  | 0.7-0.8 | 80 |
|  | 0.8-0.9 | 90 |
|  | 0.9-1 | 100 |
| **Number of entries** | 0-0.1 | 10 |
|  | 0.1-0.2 | 20 |
|  | 0.2-0.3 | 30 |
|  | 0.3-0.4 | 40 |
|  | 0.4-0.5 | 50 |
|  | 0.5-0.6 | 60 |
|  | 0.6-0.7 | 70 |
|  | 0.7-0.8 | 80 |
|  | 0.8-0.9 | 90 |
|  | 0.9-1 | 100 |

*The new anxiety scale had a maximum score of 100 points, with six indicators related to anxiety being normalized. Each 0.1 interval was recorded as 10 points, leading to a gradual increase in scores, with 100 points being the highest possible score.*

**Supplementary Table 2. Assessing the level of depression in mice.**

| **Depression indexes** | **Interval** | **Scores** |
| --- | --- | --- |
| **Sucrose preference (%)** | 0-0.1 | 10 |
|  | 0.1-0.2 | 20 |
|  | 0.2-0.3 | 30 |
|  | 0.3-0.4 | 40 |
|  | 0.4-0.5 | 50 |
|  | 0.5-0.6 | 60 |
|  | 0.6-0.7 | 70 |
|  | 0.7-0.8 | 80 |
|  | 0.8-0.9 | 90 |
|  | 0.9-1 | 100 |
| **Number of entries in the central area** | 0-0.1 | 10 |
|  | 0.1-0.2 | 20 |
|  | 0.2-0.3 | 30 |
|  | 0.3-0.4 | 40 |
|  | 0.4-0.5 | 50 |
|  | 0.5-0.6 | 60 |
|  | 0.6-0.7 | 70 |
|  | 0.7-0.8 | 80 |
|  | 0.8-0.9 | 90 |
|  | 0.9-1 | 100 |
| **Total distance (cm)** | 0-0.1 | 10 |
|  | 0.1-0.2 | 20 |
|  | 0.2-0.3 | 30 |
|  | 0.3-0.4 | 40 |
|  | 0.4-0.5 | 50 |
|  | 0.5-0.6 | 60 |
|  | 0.6-0.7 | 70 |
|  | 0.7-0.8 | 80 |
|  | 0.8-0.9 | 90 |
|  | 0.9-1 | 100 |
| **Locomotion in closed arms (cm)** | 0-0.1 | 10 |
|  | 0.1-0.2 | 20 |
|  | 0.2-0.3 | 30 |
|  | 0.3-0.4 | 40 |
|  | 0.4-0.5 | 50 |
|  | 0.5-0.6 | 60 |
|  | 0.6-0.7 | 70 |
|  | 0.7-0.8 | 80 |
|  | 0.8-0.9 | 90 |
|  | 0.9-1 | 100 |
| **Struggling time of TST (sec)** | 0-0.1 | 10 |
|  | 0.1-0.2 | 20 |
|  | 0.2-0.3 | 30 |
|  | 0.3-0.4 | 40 |
|  | 0.4-0.5 | 50 |
|  | 0.5-0.6 | 60 |
|  | 0.6-0.7 | 70 |
|  | 0.7-0.8 | 80 |
|  | 0.8-0.9 | 90 |
|  | 0.9-1 | 100 |
| **Struggling time of FST (sec)** | 0-0.1 | 10 |
|  | 0.1-0.2 | 20 |
|  | 0.2-0.3 | 30 |
|  | 0.3-0.4 | 40 |
|  | 0.4-0.5 | 50 |
|  | 0.5-0.6 | 60 |
|  | 0.6-0.7 | 70 |
|  | 0.7-0.8 | 80 |
|  | 0.8-0.9 | 90 |
|  | 0.9-1 | 100 |

*The new depression scale had a maximum score of 100 points, with six indicators related to depression being normalized. Each 0.1 interval was recorded as 10 points, resulting in a gradual increase in scores, with 100 points indicating the highest possible score.*
